# Dramatic consequences of reducing erythrocyte membrane cholesterol on *Plasmodium falciparum*

**DOI:** 10.1101/2021.10.06.463412

**Authors:** Avantika I. Ahiya, Suyash Bhatnagar, Joanne Morrisey, Josh R. Beck, Akhil B. Vaidya

**Author notes:** To whom correspondence should be addressed at Center for Molecular Parasitology, Dept. of Microbiology and Immunology, Drexel University College of Medicine, 2900 Queen Lane, Philadelphia, PA 19129.

## Abstract

Cholesterol is the most abundant lipid in the erythrocyte. During its blood stage development, the malaria parasite establishes an active cholesterol gradient across the various membrane systems within the infected erythrocyte. Interestingly, some antimalarial compounds have recently been shown to disrupt cholesterol homeostasis in intraerythrocytic stages of *Plasmodium falciparum*. These studies point to the importance of cholesterol for parasite growth. Previously, reduction of cholesterol from the erythrocyte membrane by treatment with methyl-ß-cyclodextrin (MßCD) was shown to inhibit parasite invasion and growth. In addition, MßCD treatment of trophozoite stage *P. falciparum* was shown to result in parasite expulsion from the host cell. We have revisited these phenomena by using live video microscopy, ultrastructural analysis, and response to antimalarial compounds. By using time-lapse video microscopy of fluorescently tagged parasites, we show that MßCD treatment for just 30 min causes dramatic expulsion of the trophozoite stage parasites. This forceful expulsion occurs within 10 sec. Remarkably, the plasma membrane of the host cell from which the parasite has been expelled does not appear to be compromised. The parasitophorous vacuolar membrane (PVM) continued to surround the extruded parasite, but the PVM appeared damaged. Treatment with antimalarial compounds targeting PfATP4 or PfNCR1 prevented MßCD-mediated extrusion of the parasites, pointing to a potential role of cholesterol dynamics underlying the expulsion phenomena. We also confirmed the essential role of erythrocyte plasma membrane cholesterol for invasion and growth of *P. falciparum*. This defect can be partially complemented by cholesterol and desmosterol but not with epicholesterol, revealing stereospecificity underlying cholesterol function. Overall, our studies advance previous observations and reveal unusual cell biological features underlying cholesterol depletion of infected erythrocyte plasma membrane.

**Importance:** Malaria remains a major challenge in much of the world. Symptoms of malaria are caused by the growth of parasites belonging to *Plasmodium* spp. inside the red blood cells (RBC), leading to their destruction. The parasite depends upon its host for much of its nutritional need. Cholesterol is a major lipid in the RBC plasma membrane, which is the only source of this lipid for malarial parasites. We have previously shown that certain new antimalarial compounds disrupt cholesterol homeostasis in *P. falciparum*. Here we use live time-lapse video microscopy to show dramatic expulsion of the parasite from the host RBC when the cholesterol content of the RBC is reduced. Remarkably, this expulsion is inhibited by the antimalarials that disrupt lipid homeostasis. We also show stereospecificity of cholesterol in supporting parasite growth inside RBC. Overall, these results point to a critical role of cholesterol in physiology of malaria parasites.

## Introduction

A salient feature of malaria parasites is their dependence on the host to fulfil their nutrient requirements. In addition to various nutrients, *Plasmodium* salvages lipids and fatty acids from the host. While the parasite possesses pathways for synthesizing and modifying lipids (1, 2), it lacks the machinery for *de novo* cholesterol synthesis (3-6). Unlike the intrahepatic stages where the parasites have access to copious amounts of cholesterol (7), blood stage parasites can only access cholesterol present in the erythrocyte membrane (8). Recent studies from our laboratory have identified antimalarials that inhibit two parasite plasma membrane (PPM) transporters, PfATP4 and PfNCR1, which disrupt cholesterol and lipid homeostasis in the PPM (9, 10). Upon treatment with these compounds, there is a rapid accumulation of cholesterol in the PPM rendering the parasites sensitive to the cholesterol dependent detergent saponin. This was observed as the loss of cytosolic proteins in drug treated parasites exposed to saponin. This effect was reversible indicating an active mechanism of cholesterol dynamics within the parasite (9). These unexpected consequences of exposure to novel antimalarials suggest the presence of mechanisms that influence cholesterol transport within intraerythrocytic *P. falciparum*. Previously, it was shown that intra-erythrocytic *P. falciparum* growth requires a supply of fatty acids that cannot be substituted with lipids and cholesterol from serum derived lipoproteins (11). Furthermore, studies from the Haldar laboratory have shown that cholesterol in the erythrocyte plasma membrane is essential for invasion and growth of *P. falciparum*. Interestingly, reduction of cholesterol from the erythrocyte plasma membrane by treatment with MßCD resulted in extrusion of the trophozoite stage parasites (12).

Given the observations regarding effects of new antimalarials on cholesterol dynamics and the consequences of cholesterol reduction from infected erythrocyte plasma membrane, we wished to examine a potential link between these phenomena. We have used live fluorescence video microscopy, transmission electron microscopy and treatment with antimalarials to revisit previous findings regarding the consequences of MßCD treatment of intra erythrocytic *P. falciparum*. Results from our experiments confirm and extend these observations and raise important mechanistic questions regarding the role of cholesterol in *P. falciparum* biology.

## Results

### *Dramatic expulsion of trophozoite stage parasites upon treatment with* MβCD

A previous study showed that treatment of trophozoite stage parasites with MβCD releases the parasites from the erythrocyte. In this study, the authors used filipin and ethidium bromide staining of glutaraldehyde-fixed parasites and did not assess disposition of PVM and PPM in extruded parasites (13). We aimed to examine this effect in real-time using time lapse video microscopy as well as to assess the disposition of PVM and PPM. For this purpose, we used two different transgenic *P. falciparum* lines: in one line, the gene encoding PPM-localized PfVP1 was tagged with mNeonGreen at its endogenous locus [kindly provided by Dr. Hangjun Ke; (14)]. The second line expressed two different fluorescently tagged proteins generated by using the CRISPR/Cas9 approach as described in Fig. 1 A and B. The gene encoding PVM-localized EXP2 was tagged with mRuby3 at its endogenous location, and the gene encoding the RhopH complex protein RhopH2 was tagged with mNeonGreen (Fig. 1 A and B). As shown in Fig. 1C, tagging these proteins with fluorescent markers did not affect their proper expression and localizations. EXP2 was expressed all stages and localized to the PVM, whereas RhopH2 was expressed at the highest level in mature stages of the parasite and localized to the punctate rhoptries in the schizont stage. RhopH2 is initially stored in merozoite rhoptries and secreted into the parasitophorous vacuole (PV) during invasion, eventually trafficking to the host cell membrane (15), as seen in ring stages (Fig. 1C). The use of these parasite lines permitted us to monitor the PPM, PVM and rhoptries of the parasite when exposed to MßCD using live fluorescent microscopy. The trophozoites and early schizont stage-infected erythrocytes were attached to poly-L-lysine coated plates, stained with fluorescent probes for the RBC membrane and nuclei followed by treatment with MßCD for 30 mins. MßCD was removed by washing with the culture medium and the parasites were observed by live time-lapse fluorescent video microscopy. As shown in Fig. 2A and Supplementary Video 1, the parasite extruded out of the host erythrocyte in a dramatic fashion. This expulsion occurred 10-15 min after MβCD was washed off, but the transition from being intracellular to being extracellular occurred in less than 10 seconds. The extruded parasites were surrounded by PPM (in green, Fig. 2B) as well as the PVM (in red, Fig. 2C; Supplementary Fig. 1). Depending upon the plane from which the parasite emerges from the erythrocyte, parts of the PVM were observed still tethered to the host cell. In contrast to the late stage parasites, cholesterol depletion of ring stage parasites did not result in extrusion; RhopH2 was partially localized to the erythrocyte plasma membrane in the ring stage parasites, but the parasite remained inside the host cell (Fig. 2D, Supplementary video 3). Quantification of this phenomenon showed that about 70% of the late stage parasites extruded upon MßCD treatment (Fig. 2E).

**Figure 1:**
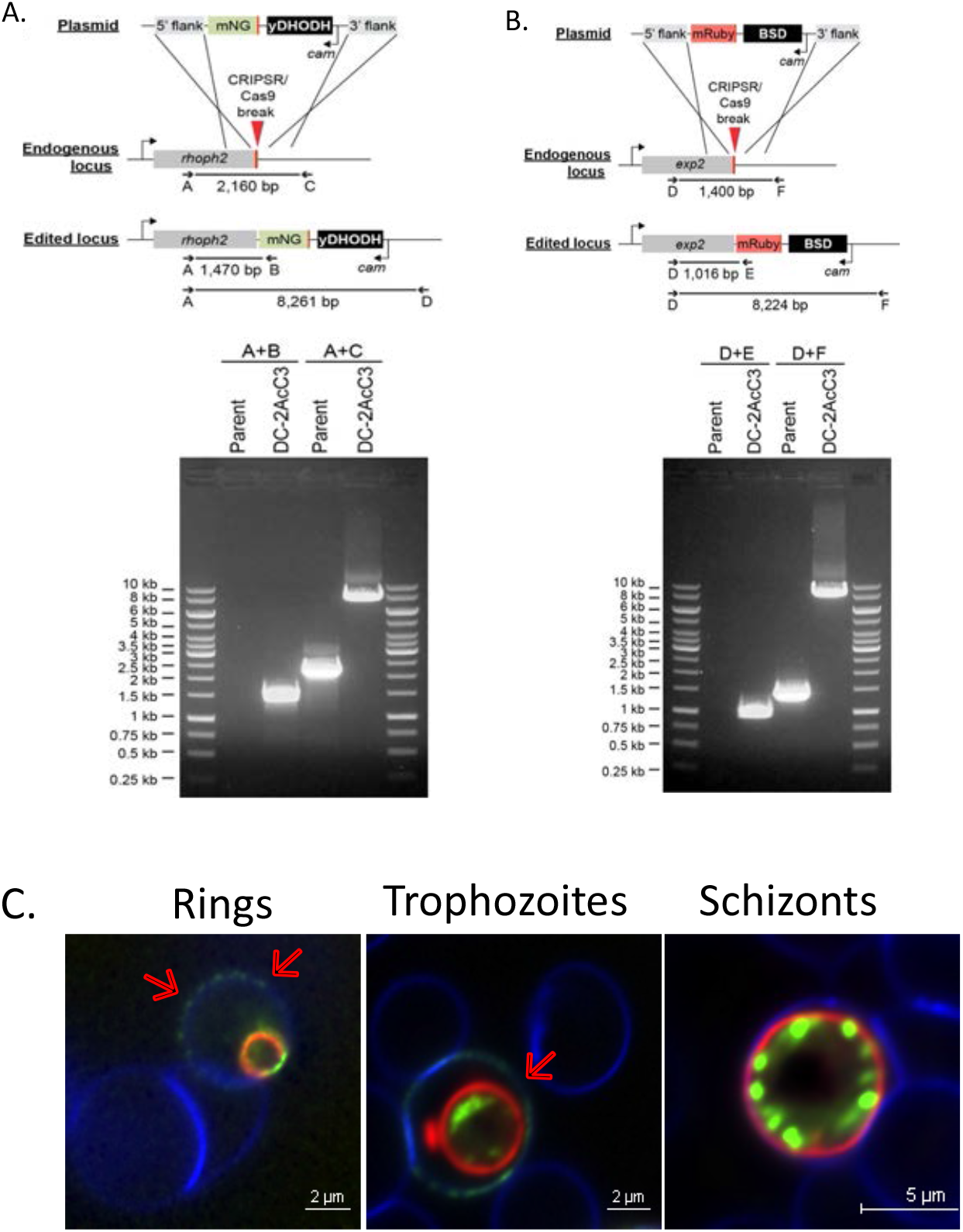
Generation of the NF54 Rhoph2/EXP2 line : (A) Strategy for tagging *rhoph2* with mNeongreen (mNG) and gel image indicating presence of clonal population expressing *rhoph2-mNG*. (B) Strategy for the tagging *exp2 with mRuby tag* and agarose gel image indicating the presence of clonal population expressing *exp2-mRuby*. Sequences of the diagnostic primers A,B,C, and D are provided in the Materials and Methods. (C) Live fluorescence microscopy images indicating the expected localization of the tagged proteins. Red arrows indicate the dual localization of RhopH2-mNG on the RBC membrane as well as the parasite in the rings and trophozoites while in the schizonts the protein is localized to rhoptries. The EXP2-mRuby marks the PVM of the parasite. Erythrocyte membrane is stained with WGA-Alexa-350 (Blue)

**Figure 2:**
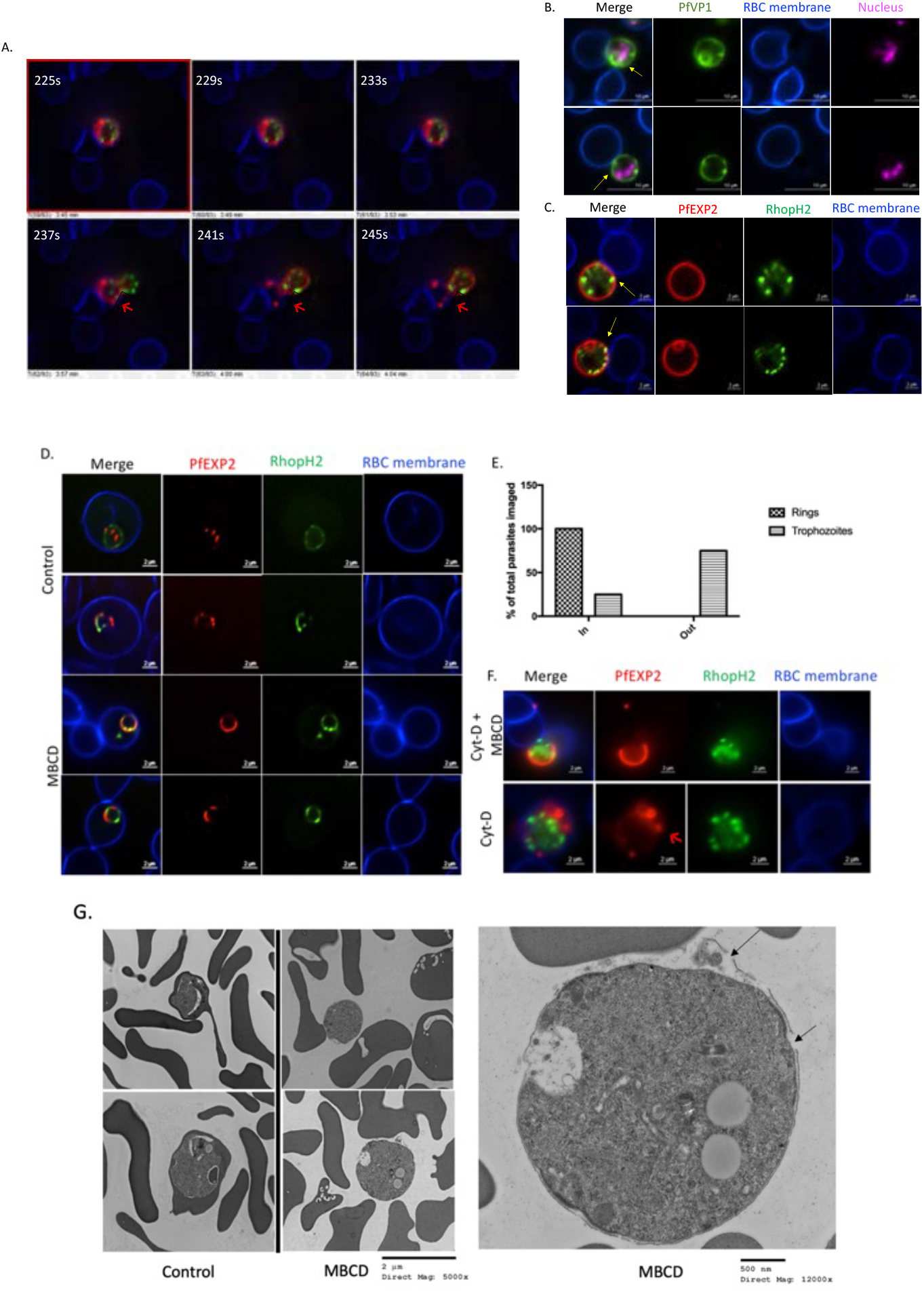
Trophozoites extrude out from erythrocyte upon treatment with MßCD. (A) Still frames from Supplementary video 1 from time-lapse video microscopy performed at. Parasites extrude out from the erythrocyte with the PVM still tethered to the erythrocyte membrane without causing the lysis of erythrocyte (indicated by arrows). (B) Representative images (>50) of PfVP1-mNG line after MßCD treatment. Nucleus is stained with SYTO-deep red (Pink). (C) Representative images (>50) of NF54 RhopH2/Exp2 line after MßCD treatment. Erythrocyte membrane is stained with WGA-Alexa-350 (Blue). Infected erythrocytes were subjected to live fluorescent microscopy after treatment with MßCD. Parasites extrude out from erythrocyte without causing lysis of the erythrocyte with their PPM (B) and PVM (C) still attached to the erythrocyte as indicated by the arrows. (D) Representative images of erythrocytes infected with ring stages of the NF54 RhopH2/Exp2 line treated with MßCD. Ring stage parasites do not extrude out from erythrocytes. (E) Quantification of parasites remaining inside/outside of erythrocytes after MßCD treatment was carried out by examining at least 100 different images of individual parasites for each experimental condition (100 for ring stages and 330 for trophozoites). (F) Upper panel: Treatment of NF54 RhopH2/Exp2 trophozoites with 0.5 μM Cyt-D for 45 mins prior to MßCD treatment does not inhibit parasite extrusion. Lower panel: Treatment with 0.5 μM Cyt-D alone does not cause extrusion of trophozoites but did alter PVM morphology (as indicated by arrows). (G) (Left)Treatment with MßCD causes extrusion of *P. falciparum* stage trophozoites without causing erythrocyte rupture unlike the control group where the trophozoite is still inside the erythrocyte (Scale bar-2μM, Direct magnification-5,000x). (Right) Zoomed in image of parasite extruded out of erythrocyte after MßCD treatment shows the PVM ruptured at multiple places indicated by black arrows. Electron microscopy of extruded *P. falciparum*. Left panels show lower magnification views of control and MβCD-treated trophozoites. A higher magnification view of an extruded parasite (right panel) shows that the PVM is compromised at multiple places (arrows) (Scale bar-500nM, Direct magnification-12,000x).

The exclusion phenomenon was also observed by phase contrast microscopy of the infected erythrocytes treated with 5mM MßCD, which also showed that the late stages extrude out of the erythrocyte without causing lysis of the erythrocytes (Supplementary Fig. 2). We assessed integrity of the erythrocyte membrane by staining MβCD treated cultures with fluorescent phalloidin. Phalloidin stains F-actin underlying the erythrocyte membrane when able to enter the cell. Parasite cultures not exposed to MβCD showed minimal phalloidin staining of either infected or uninfected erythrocytes. On the other hand, 60-70% of both uninfected and infected erythrocyte cytoskeletons were stained with phalloidin in MβCD-treated culture (Supplementary Fig. 3). These observations suggest that MβCD causes a breach in the erythrocyte membrane in a manner that allows phalloidin to enter the cell but without causing catastrophic lysis that would release most of its cytosolic content. This breach in erythrocyte membrane was seen in ring stage infected cells as well, yet this did not result in their expulsion.

The forceful expulsion of the parasite from the host cell remains mechanistically unexplained. MßCD treatment of mammalian cells has been shown to affect actin polymerization (16). Thus, one possibility could be that rearrangement and polymerization of RBC cytoskeletal actin may underlie the parasite extrusion. To test this, we treated trophozoite-infected erythrocytes with the actin polymerization inhibitor, Cytochalasin D (CytD) (17). Treatment with CytD prior to MßCD treatment did not inhibit MßCD mediated extrusion of the parasites. Treatment with CytD alone did not cause the parasites to extrude out; however, we did observe defects in the PVM of the CytD treated parasites (Fig. 2F, Supplementary video 2). Although CytD did affect the morphology of the PVM, it failed to prevent extrusion of the parasites when exposed to MßCD.

To assess the disposition of membranes of extruded parasites in some detail, we performed transmission electron microscopy of mature stage parasites treated with MßCD. The extruded parasites were still surrounded by PVM, but the PVM was compromised at multiple places (Fig. 2G). The PPM on the other hand remained intact with minimal changes in the intracellular structures. Images shown here are representative of multiple electron micrographs.

### Prior Treatment with novel antimalarials abrogates MßCD mediated parasite extrusion

We have previously shown that inhibition of PPM transporters, PfATP4 or PfNCR1 alter cholesterol dynamics in the PPM of late stage parasites (9, 10, 18). We therefore examined the effects of these cholesterol homeostasis disruptors on MßCD-mediated extrusion phenomenon. We treated late stage parasites for 2 hours with KAE609 (a PfATP4 inhibitor), MMV009108 (a PfNCR1 inhibitor) or chloroquine (a heme detoxification inhibitor) followed by cholesterol depletion with MßCD. Interestingly, the parasite extrusion was greatly reduced by prior exposure to PfATP4 and PfNCR1 inhibitors but not by chloroquine (Fig. 3A). Quantitation of this phenomenon showed that 80% of parasites treated with DMSO or chloroquine extruded following MßCD treatment, whereas only about 25% did so following treatment with PfATP4 and PfNCR1 inhibitors (Fig. 3B). This quantitation was also supported by observation of intracellular and extracellular parasites in Giemsa-stained thin blood smear (Fig. 3C and D). These results suggest that disruption of normal cholesterol dynamics in mature stage parasite results in inhibition of MßCD mediated extrusion of parasites.

**Figure 3 :**
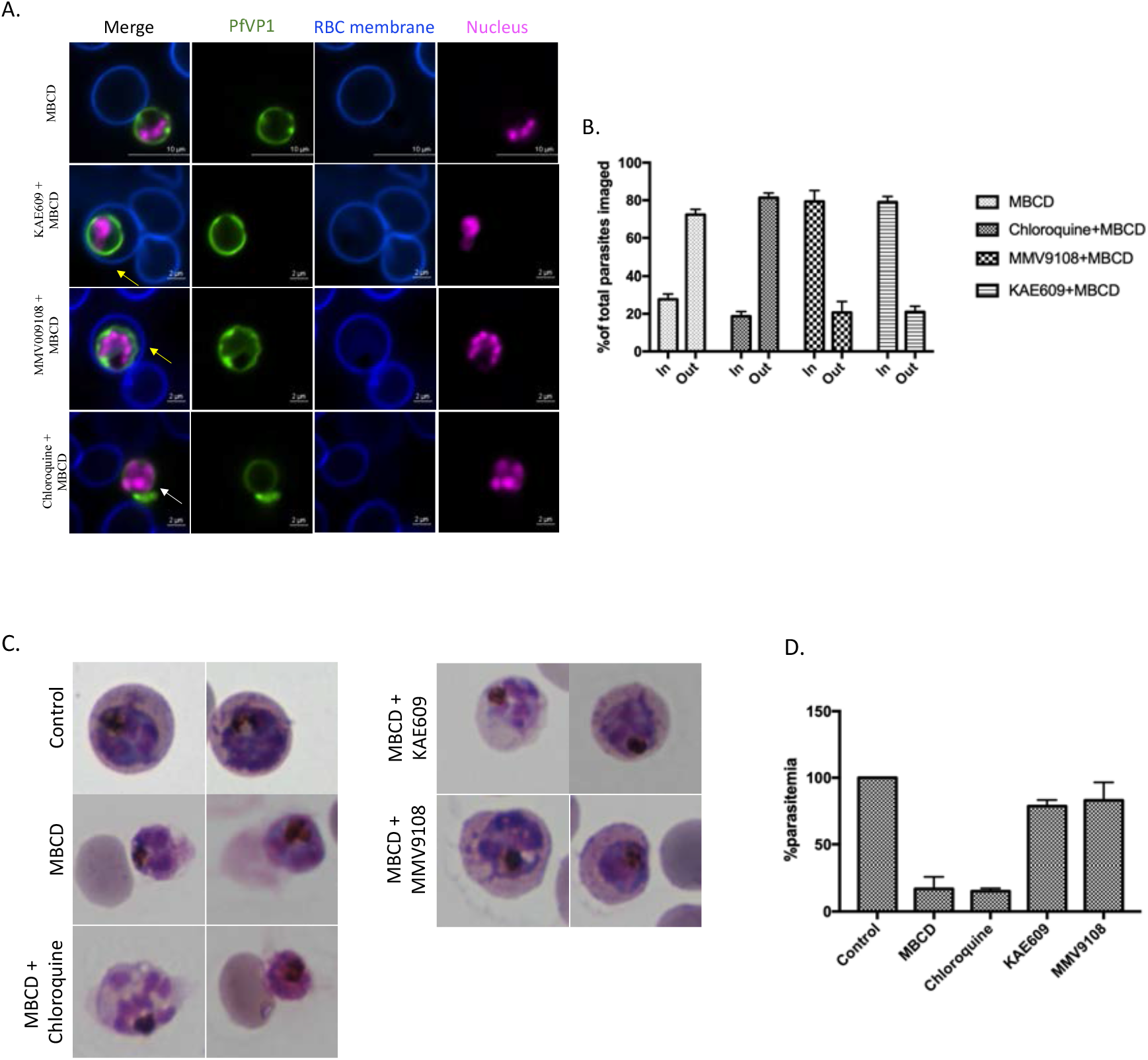
Treatment with PfATP4 or PfNCR1 inhibitors prior to MßCD treatment inhibited MßCD mediated extrusion of parasites: (A) Trophozoite stage parasites from PfVP1-mNG line were treated with KAE609 (10nM), MMV009108 (1μM) and chloroquine (150nM) prior to treatment with MßCD. Live fluorescent imaging showed that parasite extrusion was inhibited by prior treatment with KAE609 and MMV009108 (indicated by yellow arrows) while treatment with chloroquine like MßCD did not inhibit this extrusion (indicated by white arrows). Nucleus is stained with SYTO-deep red (pink). Erythrocyte membrane is stained with WGA-Alexa-350 (Blue). Representative images from 3 independent experiments for each condition (B) Quantification from live fluorescent microscopy experiments of parasite extrusion upon drug treatment followed by MßCD treatment. Approximately 250 cells for each experimental condition over 4 independent experiment were assessed. (C) Giemsa stained thin blood smears prepared from cultures after treatment with KAE609 (10nM), MMV009108 (1μM) and chloroquine (150nM) prior to treatment with MßCD. (D) Quantitation of parasite extrusion upon drug treatment followed by MßCD treatment from Giemsa-stained thin blodd smears (∼1000 cells were counted).

### Role of erythrocyte plasma membrane cholesterol in invasion and the intra-erythrocytic development of P. falciparum

Previous studies have shown that reduction of erythrocyte plasma membrane cholesterol by treatment with MßCD resulted in inhibition of parasite growth (12). Since we found that the ring stage parasites were not extruded from the erythrocyte by MßCD treatment, we went on to examine the effects of MßCD treatment of ring stage parasites for their intra-erythrocytic developmental cycle (IDC). In addition, we also examined the consequence of complementing cholesterol depleted ring stage parasites with different sterols such as cholesterol, desmosterol or epicholesterol (structures shown in Fig. 4A). We treated the ring stage infected erythrocytes with MßCD for 30 mins followed by washes with normal culture medium. Development of ring stage parasites was assessed by examination of Giemsa-stained thin smears. As shown in Fig. 4B, parasites in the control cells proceeded normally through the IDC and formed trophozoites and rings. In contrast, ring stage parasites growing in cholesterol depleted erythrocytes (MßCD treated) progressed to form what appeared to be trophozoites but failed to mature. Reconstitution with MßCD saturated with cholesterol (CD/Cho) or desmosterol (CD/Des) appeared to restore normal IDC progression. On the other hand, reconstitution with epicholesterol (CD/Epi) did not restore normal IDC, suggesting the importance of stereospecificity of cholesterol polar moiety in supporting parasite development. We also attempted to reconstitute with lanosterol or ß-sitosterol but could not assess it since these resulted in lysis of erythrocytes. To assess the viability of the ring stage parasites growing in MßCD-treated or sterol-reconstituted erythrocytes, we split the culture 1:10 into untreated normal erythrocytes. This was followed by measuring parasitemia over a subsequent four-day period. As shown in Fig 4C, ring stage parasites growing in cholesterol-depleted or epicholesterol-reconstituted cells failed to propagate over the next two cycles. On the other hand, cholesterol- or desmosterol-reconstituted ring stage parasites were able to propagate at a level of about 50% compared to the control ring stage parasites. To assess progression of ring stage parasites, we measured hypoxanthine incorporation over a 24-hour period. As shown in Fig 4D, there was significant reduction in hypoxanthine incorporation in MßCD treated and epicholesterol reconstituted parasites, whereas ring stage parasites in cholesterol or desmosterol reconstituted cells were able to incorporate hypoxanthine.

**Figure 4:**
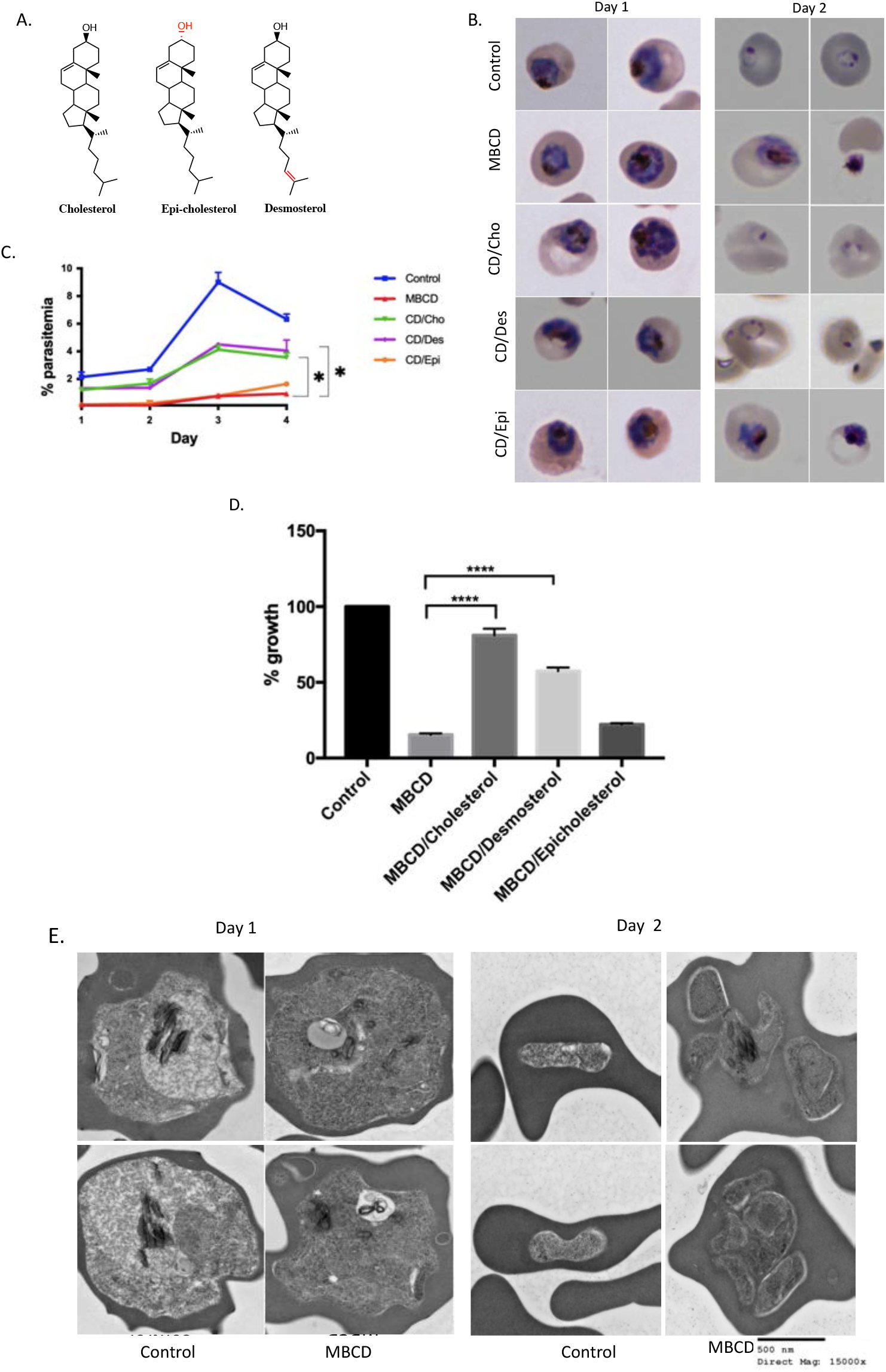
Cholesterol in the erythrocyte membrane is essential for growth and proliferation of *P. falciparum*.(A) Structure of MßCD, Cholesterol, Epicholesterol and Desmosterol. The differences between the structure of sterols are highlighted in red. (B) Giemsa-stained thin blood smears prepared from control, MßCD and erythrocytes reconstituted with cholesterol, desmosterol or epicholesterol at 24 and 48 hours after treatment. C) Parasitemia in 1:10 split of ring stage parasites from control, MßCD (CD) or erythrocytes reconstituted with cholesterol (CD/Cho), desmosterol(CD/Des) or epicholesterol (CD/Epi) over 4 days (∼3000 cells were counted). Based on one tailed unpaired *t-test* (* p<0.05). (D) Relative (^3^H)-hypoxanthine incorporation by ring stage parasites over a 24 hour period following treatment with MßCD or reconstituted with cholesterol, desmosterol and epicholesterol. Based on unpaired t-test with Welch’s correction (* p<0.05, *** p<0.001). Error bars=mean±SD N=8. (E) TEM of the ring stage erythrocyte treated with 5mM MßCD and imaged after 24 or 48 hours. (Left)After 24 hours, in erythrocyte treated with MßCD, parasites are unable to form normal trophozoites. The nuclear membrane is indistinct, hemozoin is dispersed (indicated by arrows) and food vacuole is small compared to control erythrocytes. (Right) After 48 hours, parasites in the control culture were able to invade the erythrocyte and form rings while the parasites in the MßCD treated erythrocytes undergo did not progress further and appear to disintegrate (Scale bar-500nM, Direct magnification-15,000x).

We examined the parasites by transmission electron microscopy at 24 and 48 hours in control and following MßCD treatment. As expected at 24 hours the trophozoites in the control group displayed normal morphology with distinctive internal structures and normal hemozoin formation and went on to form ring stages at 48 hours (Fig 4E, top panel; Supplementary Fig. 1). In contrast MßCD treated ring stages showed abnormal morphology with diminished food vacuoles at 24 hours. These parasites failed to develop into normal schizonts and did not egress at 48 hours after MßCD treatment. (Fig 4E, lower panel; Supplementary Fig 4). These results suggest a defect in normal trophozoite development in erythrocytes with depleted cholesterol content.

We also assessed the ability of merozoites to invade and establish infection in MßCD treated erythrocytes. MßCD treated erythrocytes were added to Percoll enriched schizonts. As shown in Fig 5A, reduced cholesterol content in uninfected erythrocytes prevented merozoite invasion. The merozoites appeared to remain attached to the surface of MßCD treated erythrocytes but failed to penetrate. Importantly, reconstitution of MßCD treated RBCs with cholesterol complemented this invasion defect (Fig. 5B). Furthermore, the parasites were able to grow in these reconstituted erythrocytes and undergo normal IDC progression. Splitting the cultures 1:10 with addition of normal erythrocytes resulted in continued normal IDC progression of parasites from control and cholesterol reconstituted cells but not from cholesterol deficient cells (Fig. 5C).

**Figure 5:**
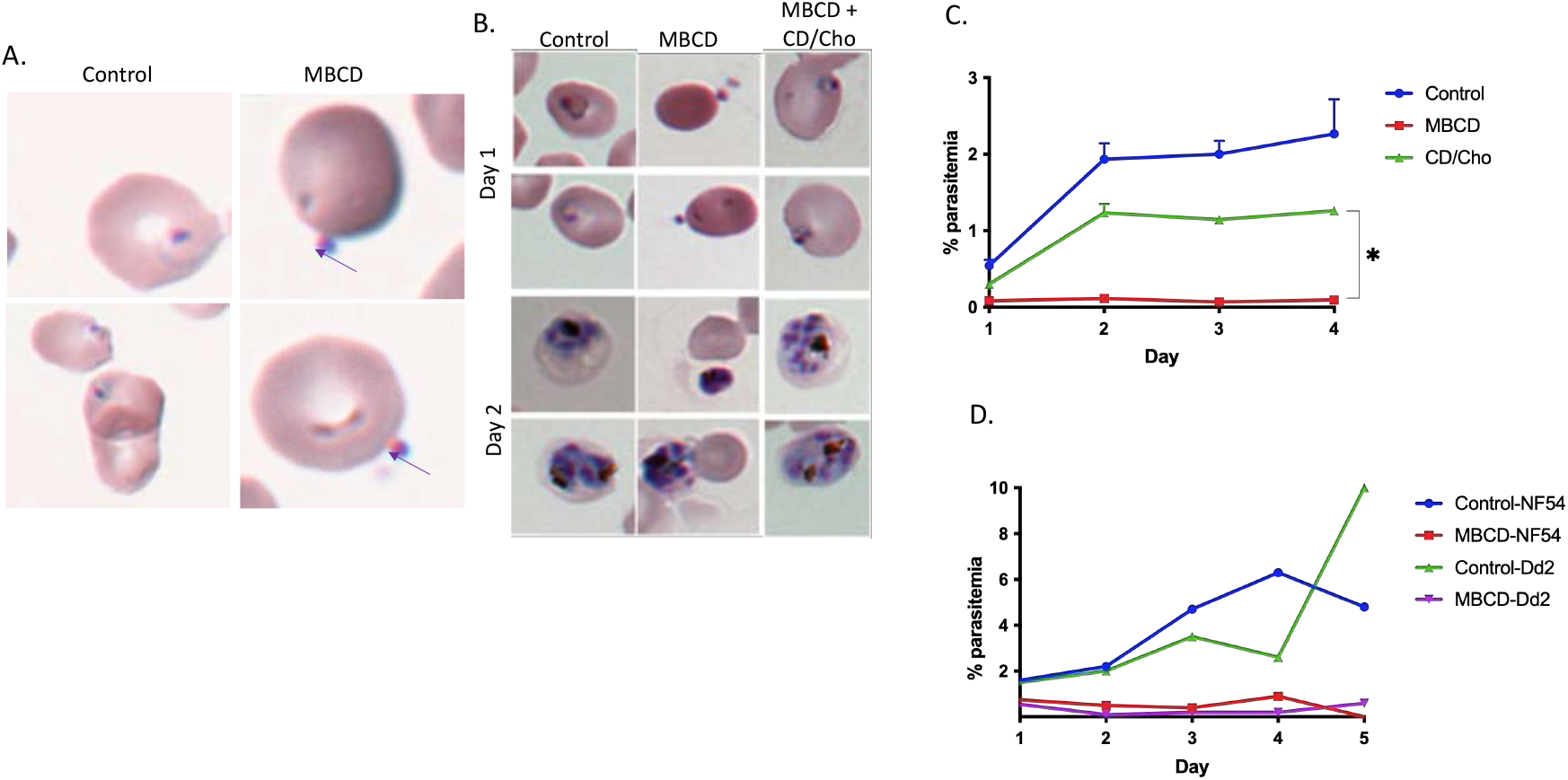
Reducing the cholesterol content in the erythrocyte membrane inhibits *P. falciparum* invasion (A) Giemsa stained thin blood smears from control and MßCD treated erythrocytes after 24 hours. While ring stages were seen in the control erythrocytes, no invasion was seen in the MßCD treated erythrocytes but merozoites appeared to remain attached to the surface as indicated by arrows. (B) Percoll enriched trophozoites were added to erythrocytes treated with MßCD or reconstituted with cholesterol and giemsa stained thin blood smears were prepared at 24 and 48 hours. Reconstituted erythrocytes were able to support parasite growth. (C) Parasitemia from 1:10 split of culture from MßCD treated and reconstituted with cholesterol in normal erythrocytes over a period of 4 days (∼3000 cells were counted). Based on Mann-Whitney test (* p<0.05) (D) Percoll synchronized trophozoites from NF54 and Dd2 strains were added onto MßCD treated erythrocytes. Parasitemia from 1:10 split of these cultures by examining Giemsa-stained thin blood smears (∼1000 cells were counted).

Different *P. falciparum* strains invade using different pathways: a sialic acid dependent or a sialic acid independent pathway (19, 20). To see if cholesterol in the erythrocyte membrane is selective for a given pathway, we added schizonts of either NF54 (sialic acid dependent) or Dd2 (sialic acid independent) strains to MßCD treated erythrocytes. Merozoites from neither strain were able to invade MßCD treated erythrocytes (Fig. 5D).

## Discussion

Using live video microscopy of fluorescently tagged parasites, we have extended previous observations showing expulsion of trophozoite stage *P. falciparum* upon treatment with MßCD. We also showed that this expulsion is inhibited by new antimalarial compounds that disrupt cholesterol homeostasis in the parasite. The dramatic expulsion of the trophozoite stage parasites following MßCD treatment remains unexplained. It is hard to perceive how an intracellular parasite can escape out of the host erythrocyte without causing catastrophic lysis of the host membrane in response to reduction in the cholesterol content in the erythrocyte plasma membrane. Expulsion of the parasite occurs after 40-45 min exposure to MβCD and after MβCD is washed off. Actual expulsion, however, occurs within seconds. The force with which the expulsion appears to occur suggest an active process mediated either by the host or the parasite. Our observation that Cytochalasin D did not prevent expulsion of trophozoites indicates that actin polymerization is not required for this process. Osmotic pressure build-up through solute uptake in trophozoites resulting in their release from the host cells has been widely used for synchronization of blood stage *P. falciparum*. However, this requires incubation with high (about 0.5 M) concentration of solutes such as sorbitol. At 5 mM concentration, MßCD exposure does not pose an osmotic challenge, and at ∼1300 Da molecular weight MßCD would not be taken up by the trophozoite-infected erythrocyte through its new permeability pathway. Furthermore, sorbitol-mediated expulsion of the parasite occurs within a few minutes and is accompanied by catastrophic lysis of the infected erythrocyte. In contrast, expulsion caused by MβCD-mediated cholesterol depletion occurs after 40-45 min when MβCD is no longer present, and the infected erythrocyte does not undergo lysis. Also, the MβCD-mediated expulsion is inhibited by prior treatment with PfATP4 and PfNCR1 inhibitors. These observations seem to argue against an osmotic imbalance as the force underlying the expulsion phenomenon described here. We speculate the possibility of a hitherto unrecognized process through which the parasite might sense cholesterol content of the erythrocyte membrane. It is of interest to note that MßCD-mediated expulsion of the parasite resembles expulsion of fungal pathogens from the mammalian cells through a process termed vomocytosis wherein the expulsion does not result in lysis of the host cell (21-23).

Inhibition of the parasite expulsion by compounds that inhibit PfATP4 or PfNCR1was unexpected; this inhibition was not a result of parasite demise since chloroquine failed to do so. We have previously shown that both these inhibitors induce cholesterol accumulation in the PPM in a reversible manner (10). PfATP4 inhibition results in Na^+^ influx into the parasite and a collapse of Na^+^ gradient across the PPM. We propose that PfNCR1 requires a Na^+^ gradient to maintain cholesterol homeostasis, and its collapse by PfATP4 inhibition results in indirect inhibition of PfNCR1. The net result of both PfATP4 and PfNCR1 inhibition, therefore, is a disruption of cholesterol homeostasis in late-stage *P. falciparum*. We envision a dynamic movement of cholesterol between various membranes of infected erythrocytes. Recent investigations have also supported the notion of cholesterol gradients in infected erythrocytes (24-28). Reduction of the plasma membrane cholesterol content, thus, appears to be sensed by the parasite in a manner that results in the expulsion of trophozoites.

The erythrocyte plasma membrane contains large amounts of cholesterol (50 mole% of total lipids). Upon infection by *Plasmodium*, the PVM acquires cholesterol but the PPM remains highly deficient in cholesterol. Indeed, this fact is the basis for widely used saponin mediated freeing of intra-erythrocytic parasites. Previous studies have suggested the importance of cholesterol for *P. falciparum* growth and invasion (12, 13, 25). A recent study using lattice light scattering microscopy demonstrated the importance erythrocyte membrane remodeling for the formation of the PVM, a process in which cholesterol is likely to play a significant role [11]. We observed that that ring stage parasites growing in cholesterol deficient erythrocytes fail to fully progress. Furthermore, the digestive vacuoles of trophozoite-like parasites at 24 hours after MßCD treatment of ring stages had unusual morphology (Fig.3E and Supplementary figure 1). At 48 hours after MßCD treatment of ring stages, parasites appeared to attempt segmentation without being able to form mature merozoites (Fig. 3E and Supplementary figure 1). One possible explanation could be that cholesterol dynamics assist lipid and/or fatty acid import to sustain parasite growth. The fact that cholesterol and desmosterol (but not epicholesterol) could substantially rescue these defects suggest stereospecificity in the role of cholesterol to sustain parasite growth. Further studies will be required to gain mechanistic insights into cholesterol dynamics and their contribution to parasite growth.

## Methods

### Parasite lines and culture conditions

Experiments were carried out using different *P. falciparum* tagged lines. 1) the 3D7 strain in which endogenous PfVP1 gene was tagged with mNeon green (mNG) at the C-terminal end (PfVP1-mNG) (14). This line was used for monitoring the dynamics of the parasite plasma membrane; 2) the NF54 strain in which endogenous RhopH2 gene was tagged with mNG and the endogenous Exp2 gene tagged with mRuby3 (NF54 Rhoph2/Exp2). Cloning was carried out with Infusion (Clontech) and NEBuilder HiFi (NEB). To generate an endogenous EXP2-mRuby3 fusion, 5’ and 3’ homology flanks targeting the 3’ end of *exp2* were PCR amplified from plasmid pyPM2GT-EXP2-mNG (29) using primers CACTATAGAACTCGAGGGAGAAACAATCTTTTATATAAAATGTACAGAGTTTGA AAG and TCCTCCACTTCCCCTAGGTTCTTTATTTTCATCTTTTTTTTCATTTTTAAATAAATC TCCAC and inserted between *XhoI* and *AvrII* sites in the plasmid pbPM2GT (30). The mRuby3 coding sequence was then amplified from plasmid pLN-HSP101-SP-mRuby3(31) using primers GATGAAAATAAAGAACCTAGGGGAAGTGGAGGAGTG and TAACTCGACGCGGCCGTCACTTGTACAGCTCGTCCATGCC and inserted between *XhoI* and *AvrII*, resulting in the plasmid pbEXP2-mRuby3. This plasmid was linearized at the *AflII* site between the 3′ and 5′ homology flanks and co-transfected with pUF-Cas9-EXP2-CT-gRNA into NF54^attB^::HSP101^DDD^ (31). Parasites were maintained in 10 μM trimethoprim (TMP) to stabilize the HSP101^DDD^ fusion and selection with 2.5 μg/ml Blasticidin S was applied 24 hours post transfection. A clonal line bearing the EXP2-mRuby3 fusion was derived by limiting dilution after parasite returned from selection and designated NF54^attB^::HSP101^DDD^+EXP2-mRuby3.

For generation of an endogenous RhopH2-mNG fusion, a gRNA target site was chosen upstream of the *rhoph2* stop codon (TCTTCACTGATTTCTTTGTA) and the gRNA seed sequence was synthesized as a sense and anti-sense primer pair (sense shown) TAAGTATATAATATTTCTTCACTGATTTCTTTGTAGTTTTAGAGCTAGAA, annealed and inserted into the *AflII* site of the plasmid pAIO3 (32), resulting in the plasmid pAIO3-RhopH2-CT-gRNA. To integrate mNG at the 3’ end of *rhoph2*, a 5’ homology flank (up to but not including the stop codon) was amplified from NF54^attB^ genomic DNA using primers AATTTCATCATTATGAAAGTTCTCAGCTTAAGAAGCATATATTAAGAATATAGTT TCAGA and CCTCCACTTCCCCTAGGACTGCTCTTCAGAATATACAGGTTTTTTATAAGATCCTCCGATATCTCCTTATATGGATCAGATATATCTGAGAAA, incorporating several synonymous mutations in the seed sequence of the gRNA target site within the *rhoph2* coding sequence. A 3’ homology flank (beginning at the endogenous stop codon) was amplified using primers GTGACACTATAGAACTCGAGTAAACGTTAAAAAAAAAATATATATAAGGAGAAA GCACTG and TCTGAAACTATATTCTTAATATATGCTTCTTAAGCTGAGAACTTTCATAATGATGA AATT, assembled in a second PCR reaction using primers GTGACACTATAGAACTCGAGTAAACGTTAAAAAAAAAATATATATAAGGAGAAA GCACTG and CCTCCACTTCCCCTAGGACTGCTCTTCAGAATATACAGGTTTTTTATAAGATCCTCCGATATCTCCTTATATGGATCAGATATATCTGAGAAA and inserted between *XhoI* and *AvrII* sites in pyPM2GT-EXP2-mNG (29) resulting in the plasmid pyPM2GT-RhopH2-mNG. This plasmid was linearized at the *AflII* site between the 3’ and 5’ homology flanks and co-transfected with pAIO3-RhopH2-CT-gRNA into NF54^attB^::HSP101^DDD^+EXP2-mRuby3. Selection with 2 μM DSM-1 was applied 24 hours post transfection (along with 10 μM trimethoprim). After parasites returned from selection, a clonal line containing the EXP2-mRuby3 and RhopH2-mNG fusions was obtained by limiting dilution and designated NF54^attB^::HSP101^DDD^+EXP2-mRuby3+RhopH2-mNG. Correct integration of the transgenes was confirmed by PCR amplifications using the following primers (indicated in Fig. 1): A:CCAGAATGTTTCGGACCATGTAC; B: TGGTATCCGGAGCCATCTACCATG; C: CACTTTGTAACTTCATTTTCTAAAATGACCTTGTTC; D: GCAACAAGTGCCTTAACCACCG; E: CGATGACTTTGATCCTCATGGTTTGC; and F: TCACTTATGTTGTATAGAGACACAATTCGT

This line was used for monitoring the parasitophorous vacuolar membrane and to mark the parasite. *P. falciparum* parasites were cultured in O^+^ human blood from Interstate Blood Bank, TN in RPMI 1640 supplemented with 2g/L sodium bicarbonate, 10mg/L hypoxanthine, 15mM HEPES, 50mg/L gentamycin sulphate, 0.5% AlbuMax II. Parasite culture was maintained at 2.5% hematocrit at 37 °C in 90% N_2_, 5% CO_2_, 5% O_2_.

### Preparation of MßCD and MßCD-sterol complexes

Parasites were treated with 5mM MßCD diluted in appropriate medium from a stock solution of 25mM MßCD (Catalog no. AC377110250) in PBS in appropriate medium. Sterols [Cholesterol (Sigma, C3045-25G), Desmosterol (Steraloids, catalog no. C3150-000), Epicholesterol (Steraloids, catalog no. C6730-000)] were loaded onto MßCD according to previously published protocol (33). Briefly, 10 μL of 15mg/ml sterols dissolved in ethanol were added to 500 μL 5% w/v MßCD, heated for 10 mins at 80°C and mixed by inverting several times until the solution was clear. The above step was repeated 4 times to add a total of 50μL sterol stock solution. The solution was heated until the sterols were stably incorporated in MßCD (clear solution). MßCD/sterol complexes were snap frozen in dry ice for 2 mins. Subsequently, the MßCD/sterol complexes were lyophilized in a speed vac until all liquid had evaporated and a fluffy powered remained at the bottom of the tube and stored at -20°C. Immediately before use, 375μl of medium was added to the MßCD/sterol complexes and vortexed until dissolved. The MßCD/sterol complexes were sterilized using 0.22μm syringe filter.

### Treatment of parasitized erythrocytes with MBCD and reconstitution with MBCD/sterol complexes

Parasite cultures were synchronized by treatment with 0.5M alanine for 10 minutes. Erythrocyte infected with ring stage parasites were treated with 5mM MßCD for 30 mins at 37 °C and washed 3 times with pre-warmed RPMI1640 media. Hematocrit was maintained at 2.5% throughout the culture and washing steps. The parasites were returned to normal culture conditions. In experiments involving reconstitution with MßCD/sterol complexes, parasites were first treated with MßCD as described above. Following this, the cells were incubated with 1:7 dilution of MßCD/sterol complexes for 30 mins. Parasites were washed 3 times and returned to normal culture conditions. Giemsa-stained thin blood smears were prepared 24 and 48 hours after the treatment with MßCD or after reconstitution with MßCD/sterol complexes. The culture at 48 hours was split 1:10 in normal erythrocytes. Growth of the parasites in these cultures were followed for 4 days by examining Giemsa stained thin blood smears

### Hypoxanthine incorporation of ring stage erythrocyte treated with MßCD or MßCD/sterol complexes

Ring stage erythrocyte (1.5% parasitemia) were treated with MßCD for 30 mins and washed 3 times with normal medium. The hematocrit was maintained at 2.5%. MßCD-treated ring stage erythrocytes were then treated with either MßCD/cholesterol, MßCD/desmosterol or MßCD/epicholesterol for 30 mins and washed 3 times with low hypoxanthine medium. The hematocrit was then adjusted to 1.5% by addition of low hypoxanthine medium to the cells. 200μl of cells were then plated in triplicates in a 96-well plate. Each well was pulsed with 22 μl of 0.5 μCi/ml ^3^H-hypoxanthine (PerkinElmer NET 177) and incubated for 24 hours at 37ºC in 5% CO_2_, 5% O_2_, 90% N_2_ chamber. Parasites were lysed by freeze/thaw and were collected on filters using a cell harvester (PerkinElmer LifeSciences). This was followed by addition of MicroScint O scintillation cocktail and incorporation of ^3^H-hypoxanthine was measured using a TopCount scintillation counter (PerkinElmer Life-sciences).

### Parasite invasion in MßCD or MßCD/Cholesterol treated erythrocytes

We treated 500 μl of 50% hematocrit erythrocytes with 5mM MßCD for 30 mins followed by 3 washes. In addition, similarly treated erythrocytes were reconstituted with MßCD/Cholesterol for 30 mins followed by 3 washes. Synchronized late stages trophozoites were enriched by centrifugation over a 70% Percoll cushion (3000 RPM, 20 mins) and washed thrice with medium. Enriched late stage trophozoites were added to MßCD-treated, MßCD/Cholesterol-reconstituted and control erythrocytes. Giemsa-stained thin blood smears were prepared 24 and 48 hours following the addition. The culture at 48 hours was split 1:10 in normal erythrocytes. Growth of the parasites in these cultures were followed for 4 days by examining Giemsa-stained thin blood smears.

### Live microscopy of parasites

Glass bottom culture dishes (35 mm) were coated with 0.1% poly-L-lysine overnight and washed 3 times with PBS. Parasite culture (250μl at 2.5% hematocrit) was added to the dishes and incubated for 30 mins to allow attachment. Culture dishes were washed 3 times with PBS followed by addition of medium. For MßCD extrusion experiments with NF54 Rhoph2/Exp2 line, all washes, incubation and imaging was done in medium supplemented with TMP. Parasites were attached to poly-L-lysine coated plates and incubated in RPMI containing wheat germ agglutinin-Alexa 350 (WGA-350) (Invitrogen, Catalog no. W7024) at 5μg/ml final concentration and incubated for 12 mins to stain the erythrocyte plasma membrane. In addition to WGA staining, PfVP1-mNG line was also stained with SYTO deep red nuclear stain for 30 min. Parasites were treated with 5mM MßCD followed by three washes with medium. The medium was replaced with phenol red free medium followed by live fluorescent microscopy. Imaging was done using the Nikon Ti microscope. WGA-Alexa-350 was visualized using DAPI filter set, mNeon green with FITC, SYTO deep red nuclear stain with Cy5 and mRuby was visualized using TRITC filter set. Video microscopy parameters are given in the legends of Supplementary Videos.

### Inhibition of MßCD mediated extrusion by novel antimalarials

To look at the effects of antimalarials on MßCD mediated extrusion, parasites were stained with WGA-350 and SYTO deep red for PfVP-1mNG line or WGA-350 for NF54 Rhoph2/Exp2 line as described above and treated with 10xEC_50_ concentrations of Choloroquine (150nM), KAE609 (10nM) or MMV009108(1μM) for 2.5 hours prior to 30 mins treatment with MßCD and washed thrice with medium. Medium supplemented with drugs (at the concentration mentioned above) was added back to the dish and live fluorescent imaging was carried out. Quantification of parasites remaining inside/outside of erythrocytes was carried out by examining at least 100 different images of individual parasites for each experimental condition.

### Cytochalasin D (Cyt-D) treatment prior to MßCD treatment of parasites

Attached NF54 RhopH2/Exp2 trophozoites were stained with WGA-350 for 12 mins, followed by three washes. The cells were then treated with 0.5μM Cyt-D for 45 mins. This was followed by the addition of 5mM MßCD for 30 mins. The culture dishes were washed 3 times and live fluorescent microscopy was performed to examine parasite extrusion. In the control condition, parasites were treated with Cyt-D without MßCD treatment.

### Transmission electron microscopy of trophozoite infected MßCD treated erythrocytes

Erythrocytes infected with trophozoite stage parasites were treated with 5mM **M**ß**CD** for 30 mins at 2.5% hematocrit followed by three washes with PBS and incubated in complete medium for 30 mins at 37 °C. Cells were washed once in PBS by gentle centrifugation and fixed with 2% paraformaldehyde, 2.5% glutaraldehyde in 100 mM cacodylate buffer and were fixed for 1 hour at room temperature. Cells were gently pelleted by centrifugation, resuspended in cacodylate buffer and immediately stored on dry ice for shipment. Ring stage infected erythrocytes were treated with 5mM for 30 mins, washed thrice with PBS and returned to culture at 37 °C. At 24 and 48 hours after the treatment, infected erythrocytes were fixed and prepared for transmission electron microscopy as described above.

### Phalloidin staining of MßCD treated cultures

Infected erythrocyte cultures from the PfVP1-mNG line were treated with 5mM MßCD for 30 mins at 37 °C and washed 3 times with prewarmed RPMI media. Phalloidin staining was carried out as described (34). Briefly, the MßCD treated cultures were stained with Phalloidin Alexa 594 (Invitrogen; diluted 1:50 from a 200 unit ml^-1^ stock in methanol) for 5 mins at 37 °C, washed thrice with prewarmed phenol red free RPMI1640. The cells were added to the glass slides and sealed with greased coverslips. Images were taken using the Nikon Ti microscope. mNeongreen was visualized using the FITC channel, Phalloidin Alexa 594 was visualized using the TRITC filter set.

## Acknowledgements

We thank the members of the Vaidya lab for lively discussion and input. We thank Dr. Hangjun Ke for providing PfVP1-mNG parasite line. This work was supported by a National Institutes of Health grant R01 AI132508 to ABV and R00 HL133453 to J.R.B.

## Supplementary Videos Legends

**Supplementary video 1:** Treatment with MßCD causes extrusion of trophozoites from erythrocytes

Representative time lapse video microscopy of the NF54 RhopH2/Exp2 line after MßCD treatment. Erythrocyte membrane is stained with WGA-Alexa-350 (Blue). Infected erythrocytes were subjected to live fluorescent microscopy after treatment with MßCD. Parasites extrude out from erythrocyte without causing lysis of the erythrocyte. Video was acquired using the Nikon Ti microscope. The interval between each frame was 4 sec and the video was captured for a total 20 min (93 frames, FPS:0.09, frames 59-64 shown in the video). Images were deconvoluted by Richardson Lucy algorithm using the Nikon NIS elements software package and converted to .avi files.

**Supplementary video 2:** Cyt-D treatment does not inhibit MßCD mediated parasite extrusion

Representative time lapse video microscopy of the NF54 RhopH2/Exp2 line after MßCD treatment. Erythrocyte membrane is stained with WGA-Alexa-350 (Blue). Infected erythrocytes were subjected to live fluorescent microscopy after treatment with Cyt-D followed by treatment with MßCD. Treatment with Cyt-D does not inhibit MßCD mediated extrusion of the parasite. Video microscopy was performed as mentioned above for video 1. The interval between each frame was 31 sec and the video was captured for a total 5 min (11 frames, FPS:0.04, all 11 frames shown in the video)

**Supplementary movie 3:** Treatment with MßCD does not cause extrusion of ring stage parasites

Erythrocytes infected with ring stages of the NF54 RhopH2/Exp2 were treated with MßCD. Erythrocyte membrane is stained with WGA-Alexa-350 (Blue). Ring stage parasites do not extrude out after MßCD treatment. Video microscopy was performed as mentioned for video 1. The interval between each frame was 31 secs and the video was captured for a total 5 mins (11 frames, FPS:0.04, all 11 frames shown in the video)

**Supplementary Figure 1:**
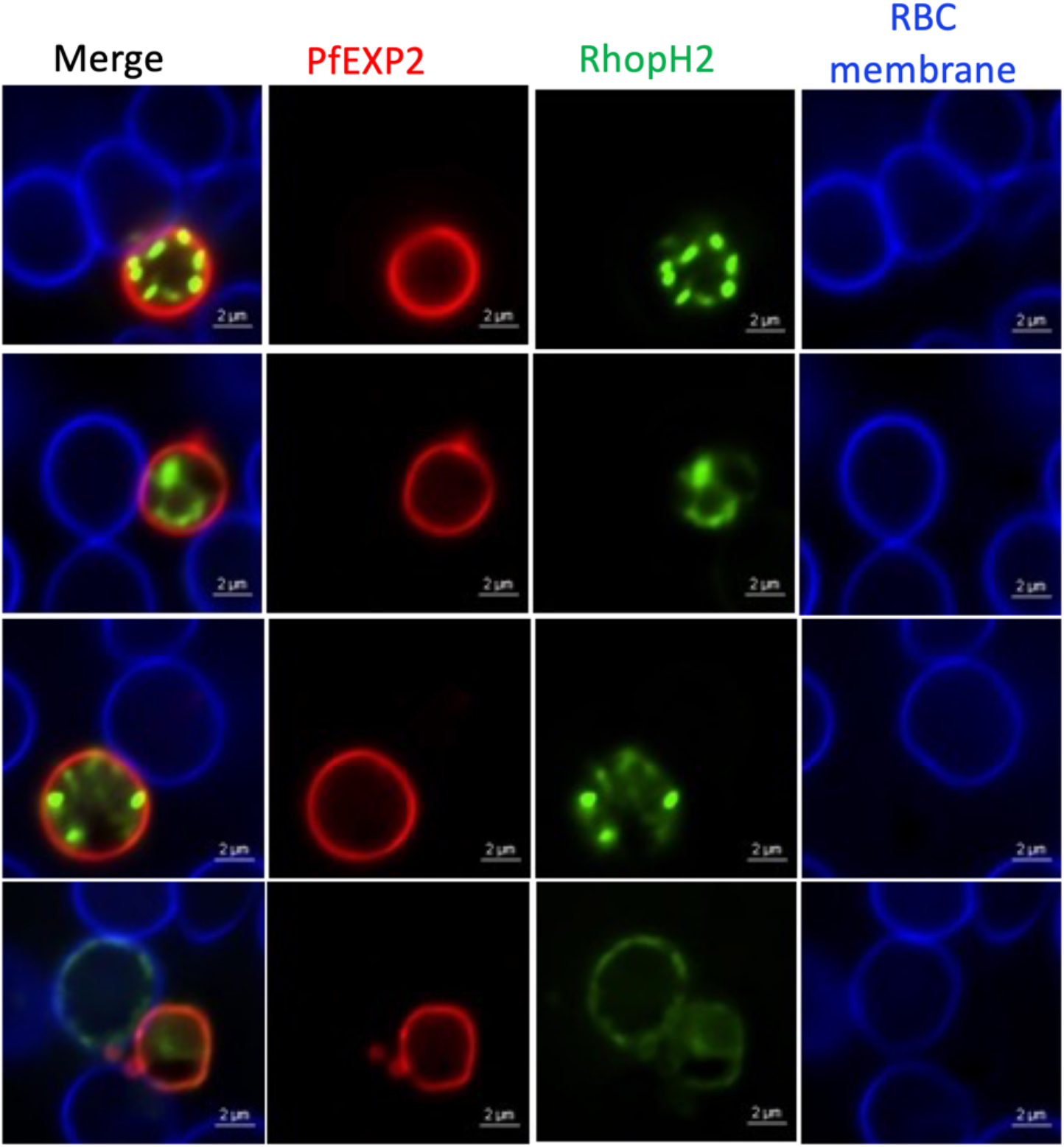
Images of extruded parasites following MßCD treatment of of the NF54 RhopH2/Exp2 transgenic line. Representative images (out of >50) of live NF54 RhopH2/Exp2 line after MßCD treatment. Erythrocyte membrane is stained with WGA-Alexa-350 (Blue), EXP2 is red and RhoH2 is green. Infected erythrocytes were subjected to live fluorescent microscopy after treatment with MßCD. Scale bar is 2 μm.

**Supplementary figure 2:**
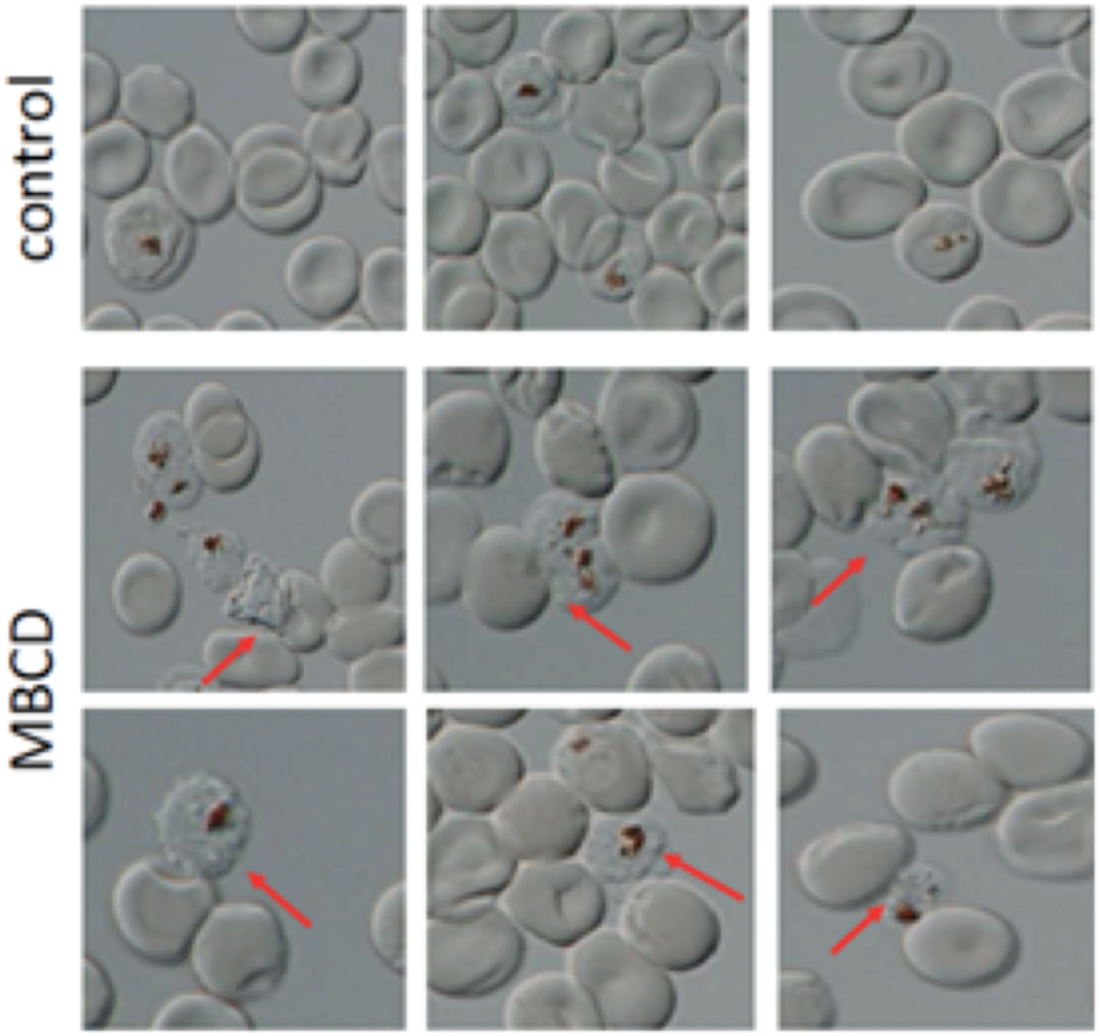
Phase contrast images of late stage *P. falciparum* extruded after treatment with MßCD. Infected erythrocytes were subjected to phase contrast microscopy after treatment with MßCD. Extruded parasites are indicated by red arrows.

**Supplementary figure 3:**
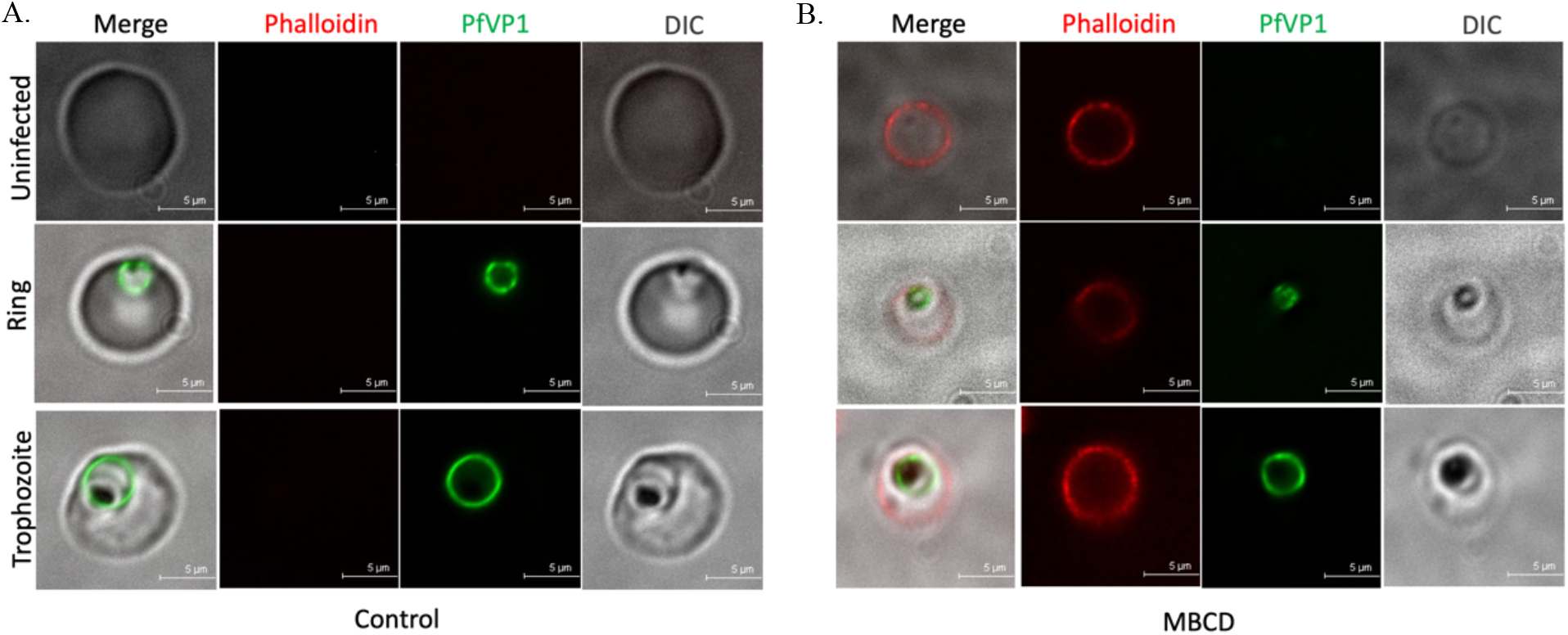
Phalloidin staining of MßCD treated cultures from PfVP1-NG transgenic line: (A) Representative images of uninfected and infected cells from untreated cultures after staining with Phalloidin AlexaFluor 594. (B) Representative images of uninfected and infected cells from MßCD treated cultures followed by staining with Phalloidin AlexaFluor 594. (Scale bar-5 μm). Overall, 60-70% of uninfected as well as infected erythrocyte cytoskeletons were stained with phalloidin, indicating membrane breach without the lysis of erythrocytes.

**Supplementary figure 4:**
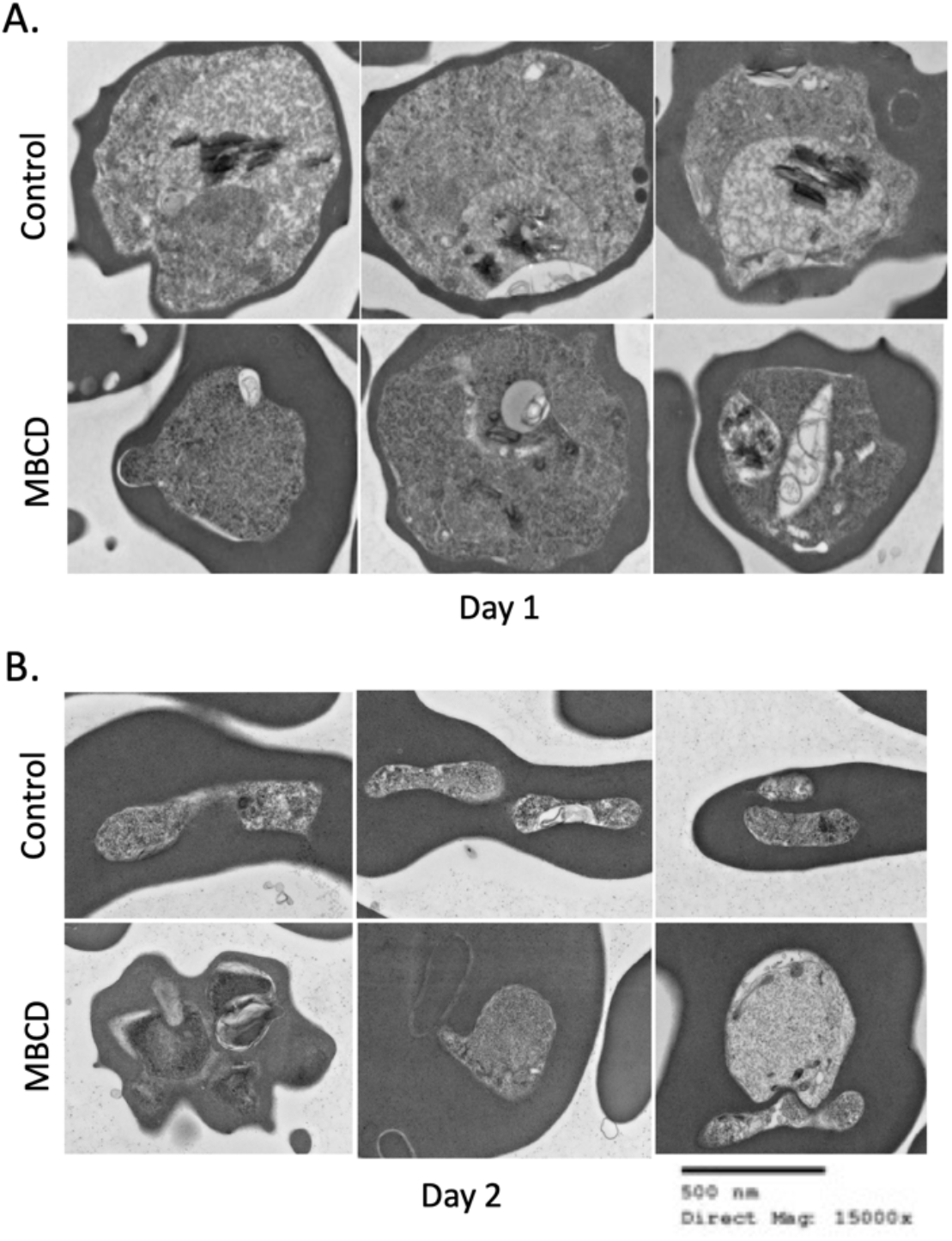
Transmission electron microscopy of the ring stage erythrocyte treated with 5mM MßCD and imaged after 24 (Day 1) or 48 hours (Day2) (A) After 24 hours, the trophozoite like parasites in the MßCD treated cultures (lower panel) have indistinct membranes, compromised food vacuole and dispersed hemozoin compared to the controls (Upper panel) (B) After 48 hours, in the control culture (Upper panel) the parasites invaded and formed rings while the parasites in the MßCD treated cultures (Lower panel) failed to form mature merozoites (Scale bar-500nM, Direct magnification-15,000x).

